# MultiCellDS: a community-developed standard for curating microenvironment-dependent multicellular data

**DOI:** 10.1101/090456

**Authors:** Samuel H. Friedman, Alexander R. A. Anderson, David M. Bortz, Alexander G. Fletcher, Hermann B. Frieboes, Ahmadreza Ghaffarizadeh, David Robert Grimes, Andrea Hawkins-Daarud, Stefan Hoehme, Edwin F. Juarez, Carl Kesselman, Roeland M.H. Merks, Shannon M. Mumenthaler, Paul K. Newton, Kerri-Ann Norton, Rishi Rawat, Russell C. Rockne, Daniel Ruderman, Jacob Scott, Suzanne S. Sindi, Jessica L. Sparks, Kristin Swanson, David B. Agus, Paul Macklin

**Author notes:** corresponding author:, www: http://MathCancer.org.

## Abstract

Exchanging and understanding scientific data and their context represents a significant barrier to advancing research, especially with respect to information siloing. Maintaining information provenance and providing data curation and quality control help overcome common concerns and barriers to the effective sharing of scientific data. To address these problems in and the unique challenges of multicellular systems, we assembled a panel composed of investigators from several disciplines to create the MultiCellular Data Standard (MultiCellDS) with a use-case driven development process. The standard includes (1) digital cell lines, which are analogous to traditional biological cell lines, to record metadata, cellular microenvironment, and cellular phenotype variables of a biological cell line, (2) digital snapshots to consistently record simulation, experimental, and clinical data for multicellular systems, and (3) collections that can logically group digital cell lines and snapshots. We have created a MultiCellular DataBase (MultiCellDB) to store digital snapshots and the 200+ digital cell lines we have generated. MultiCellDS, by having a fixed standard, enables discoverability, extensibility, maintainability, searchability, and sustainability of data, creating biological applicability and clinical utility that permits us to identify upcoming challenges to uplift biology and strategies and therapies for improving human health.

## Introduction

Over the last several years, medicine and biology have become increasingly multi-institutional and collaborative^2,11-15^. By pooling their resources, research communities can accelerate scientific progress: larger research consortia can jointly recruit patients to clinical trials that might otherwise be underpowered, create datasets that are too expensive or complex for individual institutions to build, assemble multi-parameter measurements from instruments that are too expensive or specialized to host at any one institution, and integrate diverse sets of expertise^12,14,16^.

Research communities further benefit by adopting centralized data repositories with standardized data elements^20^. Sharing data and insights is less efficient when communicating by direct member-to-member interactions with non-standard data, and “insight siloing” often results^30^. Shared repositories can spread the cost of data archiving and maintenance, while data standards make it possible to jointly develop data analysis tools. When shared repositories are publicly available, they provide a gold standard dataset promoting scientific reproducibility^31^ and the broader community can contribute further tools and secondary analyses that bring fresh expertise and insights^14,32,33^.

Recently, a consortium of 12 cancer centres in the NCI-funded Physical Sciences in Oncology Network (PSON) performed the most comprehensive characterization of two breast cell lines to date (MCF-10A: a non-tumorigenic line, and MDA-MB-231: an aggressive metastatic line)^2^. After agreeing to unified cell line sources, culturing protocols, and quality control procedures, the consortium measured the microenvironment-dependent phenotypes of the two cell lines from multiple functional points of view (e.g., cell morphology, birth rates, multiomics, genomic stability, mechanics, therapeutic resistance). This collaboration demonstrates that multi-institutional teams can study a problem in greater depth than would be possible by individual laboratories, with better cross-lab quality control and reproducibility.

However, this effort has not yet reached its full potential. As no standard exists for a systematic, standardized data representation, the measurements were shared in a collection of figures, spreadsheets, and documents, spread amongst individual and compressed files in websites^34^ and journal-hosted supplementary data^2^. This has left the data almost undiscoverable, difficult to adapt for simulation studies and other post-publication analyses, and difficult to extend by community data contributions, in contrast to centralized repositories with standardized data elements (e.g., TCGA^35^). More broadly, multicellular systems biology is in need of an extensible, orderly data format for use in cross-lab and cross-disciplinary data exchange. Otherwise, phenotypic measurements from multicellular experiments remain trapped in individual papers, and cannot readily be mined to understand the relationships between individual cell multi-omics, microenvironment-dependent cell phenotype, multicellular dynamics, and emergent disease processes that ultimately drive patient prognosis.

In this paper, we introduce MultiCellDS (MultiCellular Data Standard): a community-developed standard to functionally describe cell phenotypes with contextual information from the microenvironment. We describe the use case-driven development process, which refined the standard until it could successfully represent a library of over 200 *digital cell lines (DCLs):* a hierarchical, standardized yet extensible representation of a biological cell line’s phenotype in one or more microenvironmental contexts. Additional use cases from digital pathology and multicellular simulations drove the development of *digital snapshots*: a spatial record of the microenvironment, its cells, and their phenotypes at a single time. *Collections* allow digital snapshots and digital cell lines to be logically grouped, such as by time course (e.g., a single simulation) or study (e.g., all segmented pathology slides from a patient cohort). MultiCellDS data can include metadata to assist curation, particularly provenance (data source) when aggregating measurements from many laboratories. These standardized data elements— especially when combined with a centralised repository—can unify experimental, simulation, and clinical workflows, which often share common image and data analysis tasks, serving as biological references. See Fig. 1.

**Fig. 1:**
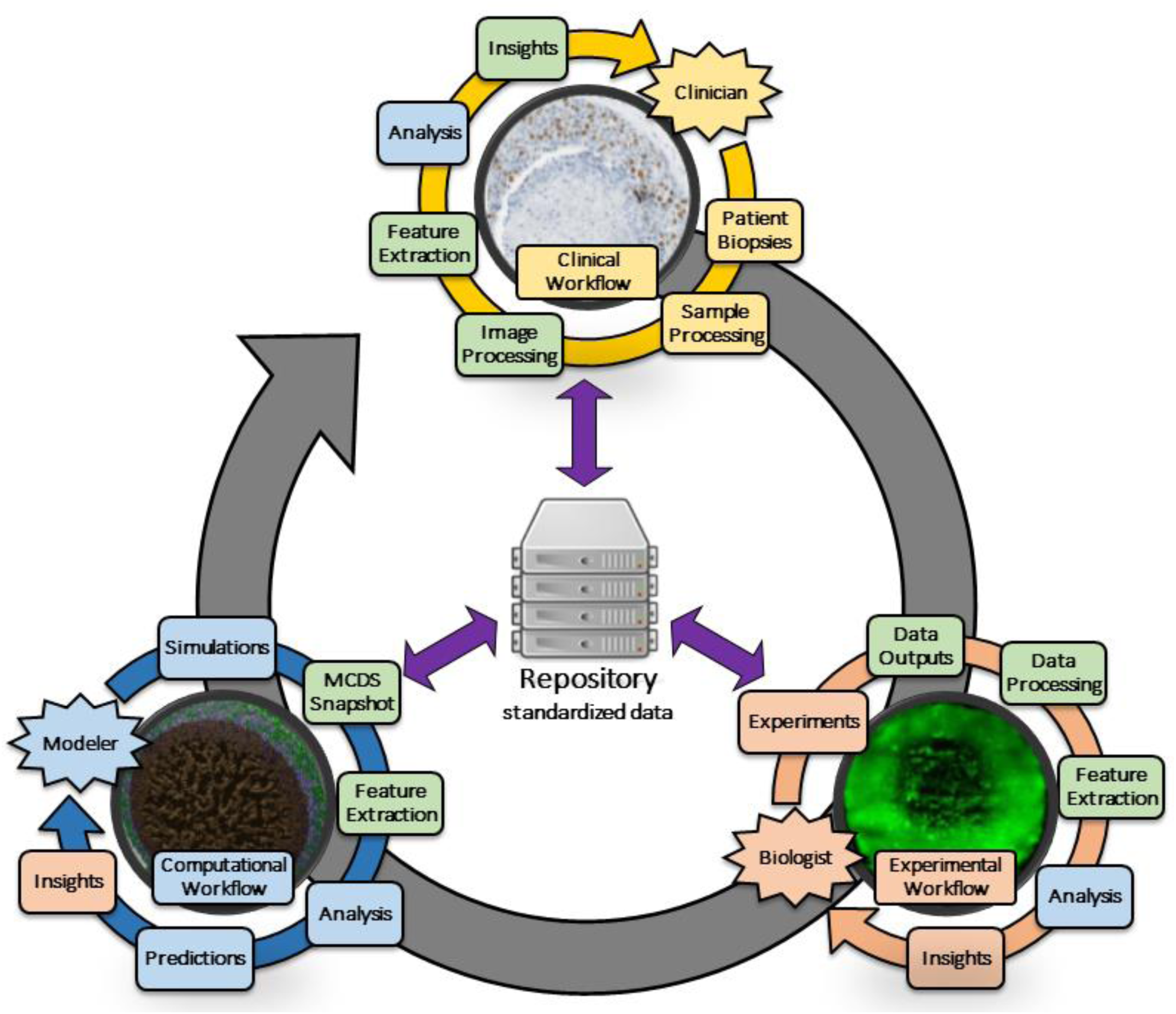
Experimental, computational, and clinical workflows all share data/image analysis (green), quantitative / analytical (blue), and biological interpretive (red) tasks. Adopting standardized data representations, along with shared data repositories, allows better integration of these workflows: standards-compliant image processing, data analysis, and machine learning software can be reused across the workflows, and computational, experimental, and clinical models can be more readily compared and cross-validated.

The MultiCellDS project is a framework for community-driven curation of biological measurements, with the goal to enable more rapid collaborative science, better quality, and enhanced reproducibility. Digital Cell Lines (DCLs) provide a standardized way of reporting what is *currently* known about context-dependent cell phenotypes and their variability, so that what is *not known* can be assessed and prioritized so as to determine the state of collective knowledge, identify the biggest gaps in that knowledge, and determine what technological advances are needed to obtain those measurements to fill in those gaps. Additionally, this project better enables researchers to investigate the impact of intra- and inter-laboratory variability in cell culturing protocols on standardized phenotype measurements, develop standardized phenotype benchmarks and reference values, improve quality control, and contribute to better reproducibility and cross-laboratory meta-analyses of experimental findings. Together with a repository to collect and share standardized, high quality data and by adopting MultiCellDS, the community has the responsibility to use, query, add, and improve the data so we can determine which cell and tissue parameters are most predictive to cell, tissue, and disease behaviour and, via a truly systematic representation of biological knowledge, rationally prioritize research directions at national and global levels, and potentially obtain the best return for limited funding resources. By collecting multiscale, multicellular data into open, public repositories makes research more reproducible, lets scientists test new biological hypotheses and therapeutic strategies, and ultimately can lay the groundwork to improve human health.

## Results

A multidisciplinary, multi-institutional team of computational modellers, biologists, clinicians, engineers, and data scientists met virtually and in person to create a standard for multicellular phenotype data via use case development. Thus, our results are both a data standard and a starting library of standardized data. (See **Method** and **Supplementary Materials** for further details.) We created three iteratively refined primary data types: a *digital cell line (DCL)*, a *digital snapshot*, and a *collections* type. We summarize these data types and their development below; full details can be found in the **Supplementary Materials** and at MultiCellDS.org.

### Data type: Digital cell lines

A *digital cell line (DCL)* is a consistent, hierarchical organization of quantitative phenotype data for a single biological cell line, including the microenvironmental context of the measurements (which further organizes the data) and essential metadata (“data about data” such as biological classification, curation, quality, and citation information; see below). Its phenotype data elements are organized by function (Table 1) to help systemize information, while allowing flexibility and extensibility through user-defined custom data elements. The standard was refined until we could successfully represent a phenotype measurements for a library of over 200 cell types, including patient-derived and “standard” cultured cancer cell lines, endothelial cells (to test the standard against mesenchymal cells), yeast (to test against non-mammalian, non-cancer lines), and bacteria (to ensure the standard could represent prokaryotic cell data).

**Table 1.**
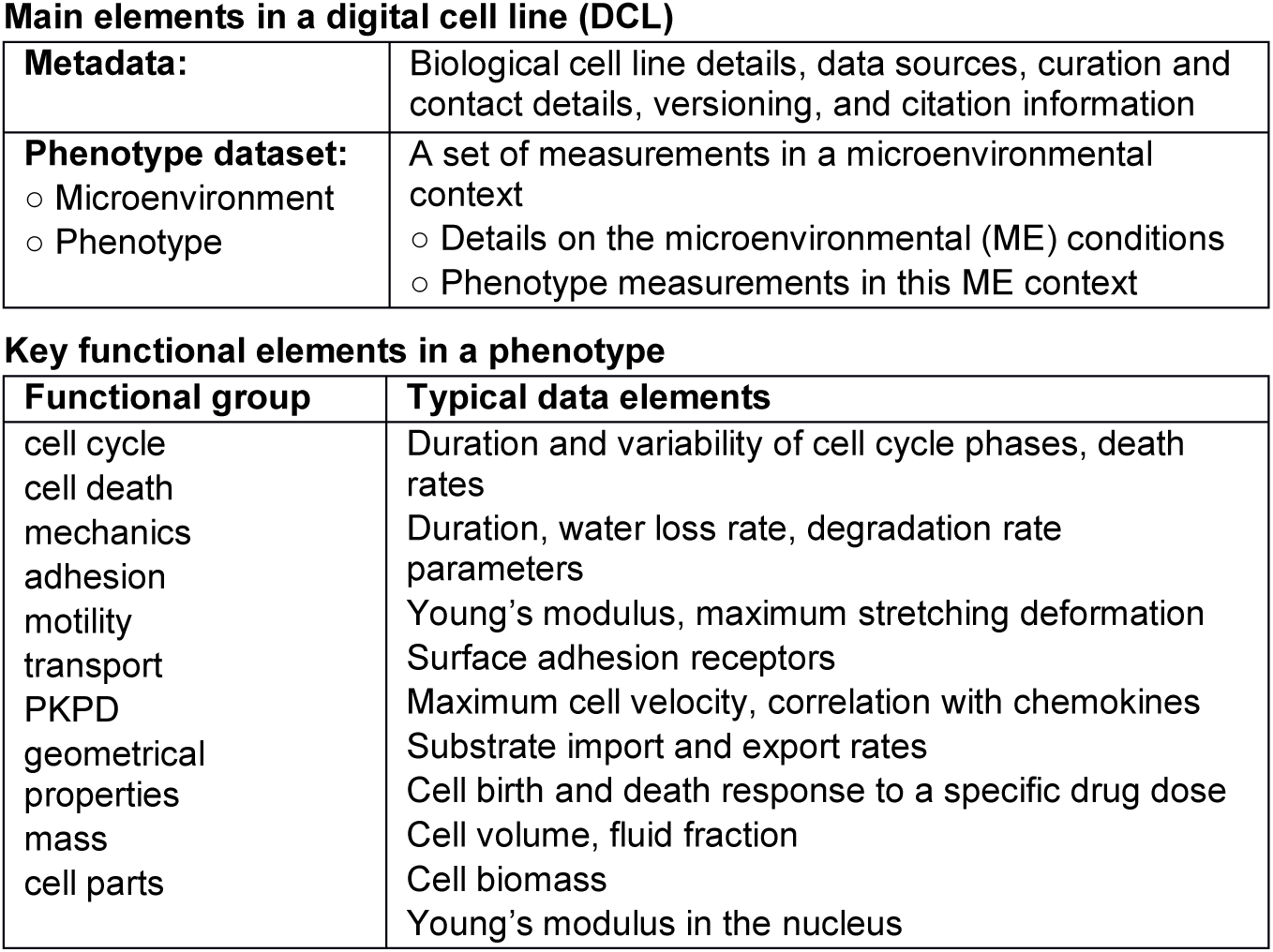
Key functional parts of a digital cell line (top) and phenotype (bottom).

Each DCL has two main parts: cell line metadata and one or more phenotype datasets. Cell line metadata include information about the personnel who created and maintain a DCL, where the data came from, and other information. A phenotype dataset functionally organizes currently available cell phenotypic measurements in a specific (and annotated) microenvironmental context for the measurements (e.g. phenotype measurements in hypoxic conditions). See Table 1.

To record the data, we use XML (eXtensible Markup Language), a hierarchical, structured ordering of data elements that can be tailored to a specific domain using XML Schemas^36,37^ and related tools. Additionally, the data can be stored in databases. XML is an ideal format for collaboratively developing MultiCellDS due to its human-editable format, the availability of many pre-existing software packages, the ecosystem of related technologies (e.g. XPATH^38^), and its widespread support across simulation and data analytics software.

#### Recording the cell phenotype

Cellular behaviour measurements are grouped hierarchically by function in the phenotype element. The main functional groups record parameters associated with the cell cycle (including phase information), cell death, mechanics, motility, adhesion, pharmacodynamics (denoted by PKPD to include pharmacokinetic drug metadata), transport processes of chemical entities, geometrical properties, and mass (see Table 1). The level of detail describing the cell cycle can vary with the experimental or clinical setup (e.g., clinical pathology often uses Ki-67 positive / negative status to quantify proliferation^39^, while experimental investigations often annotate cell cycle status as G_0_/G_1_ or S/G_2_/M (e.g., using FUCCI^40^). Thus, we allow multiple cell cycle representations in a phenotype dataset. PKPD describes the pharmacokinetic of a drug response (e.g., changes in cell proliferation or motility) and drug pharmacodynamics for individual drugs (at one or more dosages) or combinations of drugs. In the future, we plan to integrate the PKPD data elements with PharmML^41^.

#### Microenvironment: context for the phenotype

A cell’s microenvironment provides the context for its phenotype by indicating the external properties of a cell, like the mechanical stress and levels of chemical substrates in the immediate vicinity of the cell. Cells often behave differently in different microenvironments. For example, the cell birth rate decreases and the mean time spent in G_0_/G_1_ can increase for some cell lines when oxygen is reduced from normoxic (21% O_2_) conditions to hypoxic (0.1% O_2_) conditions^42,43^. The microenvironmental context for a phenotype measurement set is critical, not only for comparing data across experiments and cell types, but also for building computational models^6,44^. This observational approach allows us to link microenvironment and phenotype data without fully determining subcellular mechanisms (e.g., metabolic pathways). Since the data for these microenvironmental measurements can come from multiple instruments with different sampling resolutions, or even order-of-magnitude estimates, we permit multiple domains in the microenvironment data element. The microenvironment data element defines one or more variables (e.g., extracellular oxygen concentration), annotates them against existing ontologies like Chemical Entities of Biological Interest (ChEBI)^45^, and then records the value of each variable. See the Supplementary Information for more details.

#### Phenotype Dataset: connecting phenotype, context, and other scales

Each phenotype element is combined with a microenvironment element to form a *phenotype dataset* (a tuple of elements). This allows mapping a cell line from a position in the microenvironment space (all possible combinations of microenvironmental conditions) to a position in the phenotype space (the space of all possible phenotypes). We use keywords to denote differences between the phenotype datasets (e.g. different levels of oxygenation) to make searching between phenotype datasets easier. See the Supplementary Material for further examples. In the long term, we plan to assemble DCLs for many different phenotype datasets (breadth), with each phenotype dataset having as much multi-scale, multi-omics data as possible (depth).

#### Measurement-level metadata

Several metadata elements can be associated with any MultiCellDS measurement as XML attributes. The attached metadata will deal with measurement variability (e.g., standard deviation), uncertainty (e.g., margin of error), units (if applicable), and type. We annotate units with the Ontology of Units of Measure (OM)^46^, which allows prefixing of units and multiplication and powers of units. Measurement type allows us to indicate whether a measurement is raw (straight off an instrument), direct (directly measured by a specialised instrument), inferred (derived by fitting a mathematical model to raw or direct data), literature (extracted from published data for the same cell type and measurement conditions), estimated (estimated to order-of-magnitude based upon published data for similar cells in similar conditions), or assumed (e.g., a mathematical simplification). See the Supplementary Material for more details.

#### Cell Line Metadata: information about the digital cell line

These metadata record information about an overall DCL rather than about individual measurements. Each DCL has a unique identifier (using the standards developed at identifiers.org^47^) and versioning information to help reproducibility. We record information about the people involved with a DCL (Creator, Curator, Current Contact, and Last Modified By) using well-established ORCID^48^ elements. When measurements are mined from pre-existing studies, we record the source publications and analysis protocols. Next, we record biological or clinical information like what species, strain, and/or organ the cell came from, pathological or clinical classification, and any other unique identifying information. See the Supplementary Material for further details.

#### Provenance, Reproducibility, and Aggregated Citations

DCLs can aggregate measurements from multiple sources, using interpretations and analyses from several contributors. Over time, community-driven improvements lead to increasingly complex provenance for DCLs. The review process (see Methods) determined that DCLs must track and properly annotate this history to ensure both reproducibility and correct attribution while encouraging data contributions from the community. DCLs track data origins, creators and curators, and analysis software / algorithms as metadata. This information can be used to create an aggregated citation that properly includes this information. For example:

> “We used digital cell line CELL_LINE_NAME [R_1_,R_2_] version ## (MultiCellDB ##), created with data and contributions from [R_3_,R_4_-R_n_].”

Here, R_1_ is the publication that originally introduced the DCL, R_2_ is the publication that introduced the latest version (if different from R_1_), MultiCellDB ## is a unique identifier in the MultiCellDB repository, R_3_ is the present paper (to define the MultiCellDS data elements), and R_4_-R_n_ are publications for (1) the original data used to create the DCLs, (2) software and algorithms used to analyse the data, and (3) previous versions of the DCL between the original and current version. This form of citation emphasizes data contributors (R_4_-R_n_), data curators (R_1_,R_2_), and analytics software developers (R_4_-R_n_).

#### Digital Cell Line Numbering and Versioning

Each DCL is assigned a unique identifier to distinguish it from other DCLs, while allowing simple identification of related DCLs. See Fig. 2 for further details. We demonstrate the interplay of the branches and versions with curatorial control in Fig. 3, using concepts originating in version control software systems such as git^49^ that support forks, branches, and merges of projects with version tracking.

**Fig 2.**
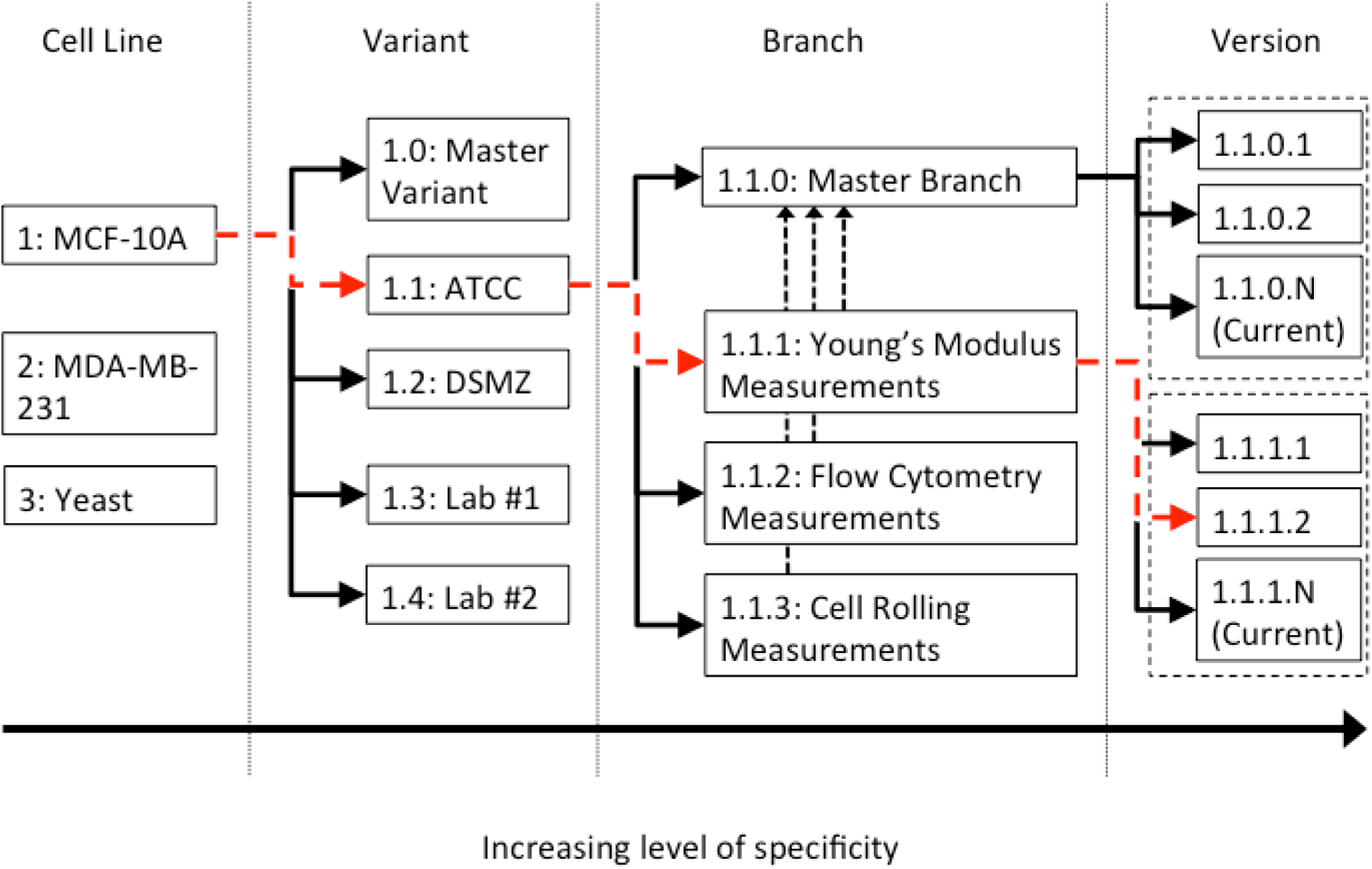
Systematic version-based numbering: Each number has four parts: Line.Variant.Branch.Version. A <u>Line</u> represents a distinct type of cells, e.g. MCF-7. Each <u>Variant</u> represents a (sub-)line associated with a particular organization or group that may have customized the cell line, e.g. ATCC or an investigator’s group. Each list of variants is independent of the parent line. The 0^th^ variant represents the collective behaviour aggregated over all variants. Each <u>Branch</u> represents a particular set of (curated or uncurated) measurements. For example, one branch could have curated Young’s Modulus measurements and another branch could have uncurated cell cycle phase durations. The 0^th^ branch is the curated version with the most data included, pending any potential conflicts in the data between various branches. The <u>Version</u> represents the improvement (increase in quantity or quality) of the data in the DCL, thus recording the history and evolution of the data. By having lines, variants, branches, and versions of DCLs, we can record new data without impacting any preexisting DCLs. In this example, the red line shows DCL 1.1.1.2 for the MCF-10A line with the ATCC variant with Young’s Modulus measurements and the second version of the data.

**Fig 3.**
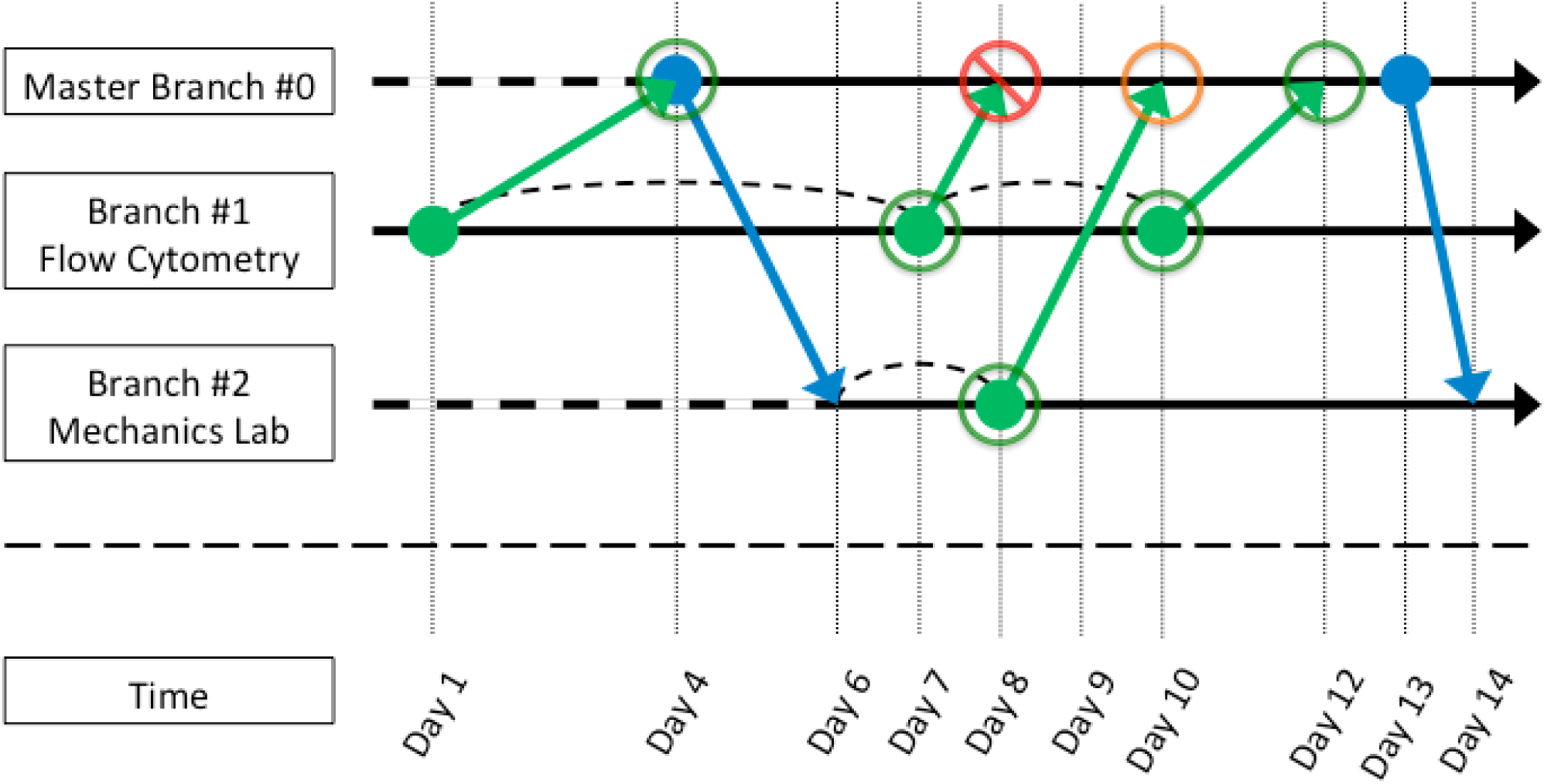
Interaction of branches and versions (example). Initially, a flow cytometry group contributes data in a new branch (#1) and merges it into the master branch (#0). A previously designated master branch curator accepts the merge. Later, the flow cytometry group mistakenly updates its branch (#1) with faulty data; the curator rejects the merge to the master branch. Meanwhile, a cell mechanics group forks (duplicates) the master branch to create a new mechanics branch (#2), adds their data to the new branch, and merges this data into the master branch. Because the master branch curator cannot evaluate the mechanics data and it does not conflict with any pre-existing data, the data are automatically accepted into the master branch. The flow cytometry group collects new data to revise their branch (#1), and the master branch curator approves the merge. Lastly, the mechanics group seeks the newly curated flow cytometry data and merges the master branch into their mechanics branch (#2). Each successful commit or merge results in a new version of the DCL. In terms of access, all data should be publically available, unless a private branch has been created (forking). We purposely limit describing any policies to let each community develop their own standards. See **Supplementary Material: Norms for MultiCellDS**. **Legend:** Solid circles are commits, green solid upward diagonal lines are merges, blue solid downward diagonal lines are pulls, unfilled circles are conflicts and merges, and curved dashed lines are the introduction of additional data.

### Data type: Digital Snapshots

DCLs record averaged cell phenotype measurements and microenvironmental context on a static basis. However, it is important to record spatial and dynamical information as well. Motivated by use cases arising from recording segmented pathology images and cancer single-time simulation outputs, we extended the data elements from DCLs to create *digital snapshots*: a digitized record of the spatial distribution and phenotypic state of all cells in a specified region at a single point in time, along with spatial microenvironmental measurements. Digital snapshots can be applied to experimental, clinical or computational data, and they can describe either individual cells, clusters of cells, or cell densities. The review panel iterated on these data elements until they could sufficiently describe data outputs for several classes of computational models^50^, including continuum models^28,29,51,52^ cellular automata^53,54^, cellular Potts models^55,56^, agent-based models^51,57^, and vascular network models^58,59^. See recent reviews for definitions of these models^51,57^. The panel further iterated the data elements to ensure that digital snapshots could sufficiently represent segmented pathology data from clinical and experimental samples^17^. See Supplementary Material **Human Overview** for further details.

### Data type: Collections

The digital snapshot uses cases also exposed the need for logically bundling related digital snapshots, such as all saved times in a single simulation experiment or a patient cohort. Thus, we created the *collections* data type, which can bundle any combination of DCLs, digital snapshots, or other collections. The groupings permit novel and unforeseen combinations to emerge as the community continues to develop ideas about how the data should be aggregated. For example, each simulation in a computational study can now be bundled as a collection of digital snapshots (a time course), along with the DCLs used in the simulation. These collections themselves could be bundled in a collection to more easily share and archive the paper’s data.

### Ontology definitions, software, and extensibility

While the XML schemas specify the structure of MultiCellDS, we must also annotate the meaning of the MultiCellDS elements, allowing a consistent definition of individual terms. To that end, we generated an OWL (Web Ontology Language)^60,61^ ontology from the XML schemas, permitting a mapping of each element onto its ontological counterpart. (See Box 1 to define these terms.) We extensively link to other ontologies (e.g. Gene Ontology (GO)^62,63^, Cell Behavior Ontology (CBO)^64^, Phenotypic Quality Ontology (PATO)^65^, Ontology of Physics for Biology (OPB)^66^), as these expert-defined ontologies provide excellent, time-tested definitions. Whenever possible, we adopted definitions from these pre-existing ontologies. Where terms were non-existent or poorly suited to multicellular systems biology, we defined new terms; these are defined in the OWL files. We used CodeSynthesis’ XSD/e^67^ and PyXB^68^ to create APIs to read and write MultiCellDS from C++ and Python respectively. We expanded the software to automatically generate OWL files from our XML schemas.

##### Box 1. Summary of digital snapshots in the library

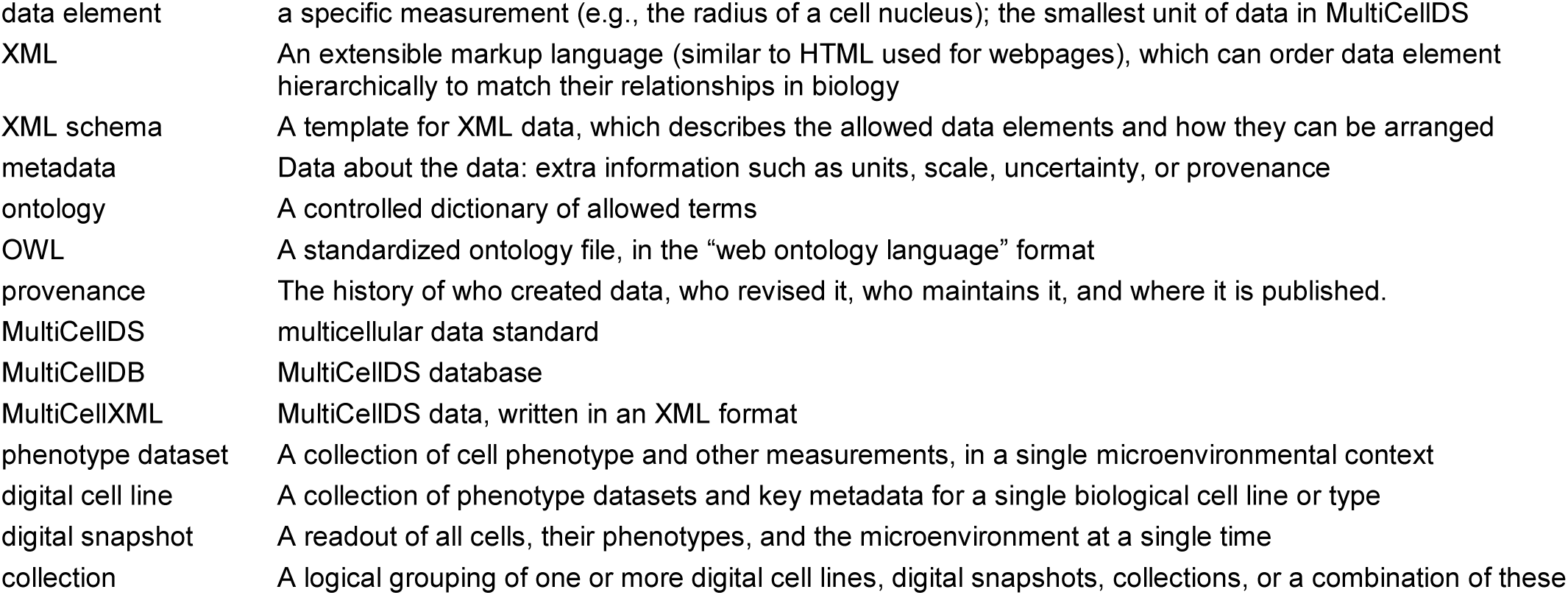

We anticipate that advances in experimental and theoretical biology and clinical science will necessitate new, descriptive data elements. To encode such data in MultiCellDS without clashing with standards-compliant software, custom data can be inserted within clearly delineated <custom> data elements anywhere in the data hierarchy. This allows scientists to rapidly adapt MultiCellDS to their needs without the potential bottleneck of requiring a full standards committee review and approval of updates to the standard. The standards panel will regularly analyse the most prevalent custom data elements for inclusion in future releases of MultiCellDS. The use of these custom elements demonstrates how MultiCellDS can seamlessly adapt to the changing needs of the research community, and how draft data elements can eventually become part of the full standard.

### Digital Cell Line and Snapshot Library: MultiCellular DataBase (MultiCellDB)

We created a public repository, the MultiCellular DataBase (MultiCellDB), at http://MultiCellDS.org/MultiCellDB, in order to store the MultiCellDS files. This repository provides web based access to digital cells lines. The repository is based on the DERIVA scientific asset management system^69^. MultiCellDB allows contributors to upload DCL files which are stored in their original form, as well as converted into an entity-relationship model for storage in a relational database. The MultiCellDB interface provides a faceted search capability and allows users to search for cell line data by phenotype, measurement value, microenvironment, or metadata value. DERIVA provides a fine grain access control mechanism which allows data sets to be quarantined while they are being curated prior to public release.

To help drive development of the standard, we developed a library of over 200 DCLs. While most of these DCLs are for human cancer, we worked to ensure that the standard could handle non-cancerous and non-epithelial human cells (HUVECs), non-human mammalian cells (murine lymphoma), single celled eukaryotes (*S. cerevisiae*), and prokaryotes (*S. aureus* and *E. coli*). See Table 2 for a complete listing and an overview of how the standard evolved due to the development of the DCLs.

**Table 2.** Summary of digital cell lines in the library.

Collections of digital cell lines
| Collection | Number | Description | New or Improved Elements |
| --- | --- | --- | --- |
| PSON | 2 | Very detailed MCF-10A and MDA-MB-231 human breast cell lines based upon PSON work.<sup>2</sup> | "Dimensions" of results, e.g. force vs. indentation amount |
| ATCC | 38 | cancer cell lines built from publicly-available ATCC data sheets<sup>5</sup> | None |
| GBM | 39 (133) | glioblastoma multiforme (aggressive brain cancer) based on patient data<sup>10</sup>. 39 detailed lines, 133 additional partial lines. | Patient and Disease Elements |
| DCIS | 17 | ductal carcinoma in situ of the breast, built from previously published patient pathology data<sup>18</sup> and modest extensions of earlier parameter estimation protocols<sup>6,19</sup>. | Patient and Disease Elements |
| endothelial | 1 | endothelial cell lines, built from prior literature<sup>16,21</sup> for HUVECs | None |
| yeast | 1 | *species* based on previously-published data<sup>22-25</sup> (first non-mammalian digital cell line) | Generation number |
| bacteria | 2 | *species* based on previously-published data<sup>26,27</sup> (first non-eukaryotic digital cell lines) | None |
| lymphoma | 2 | murine lymphoma lines based upon previously published experimental/simulation studies<sup>28,29</sup> | None |

**Table 3.**
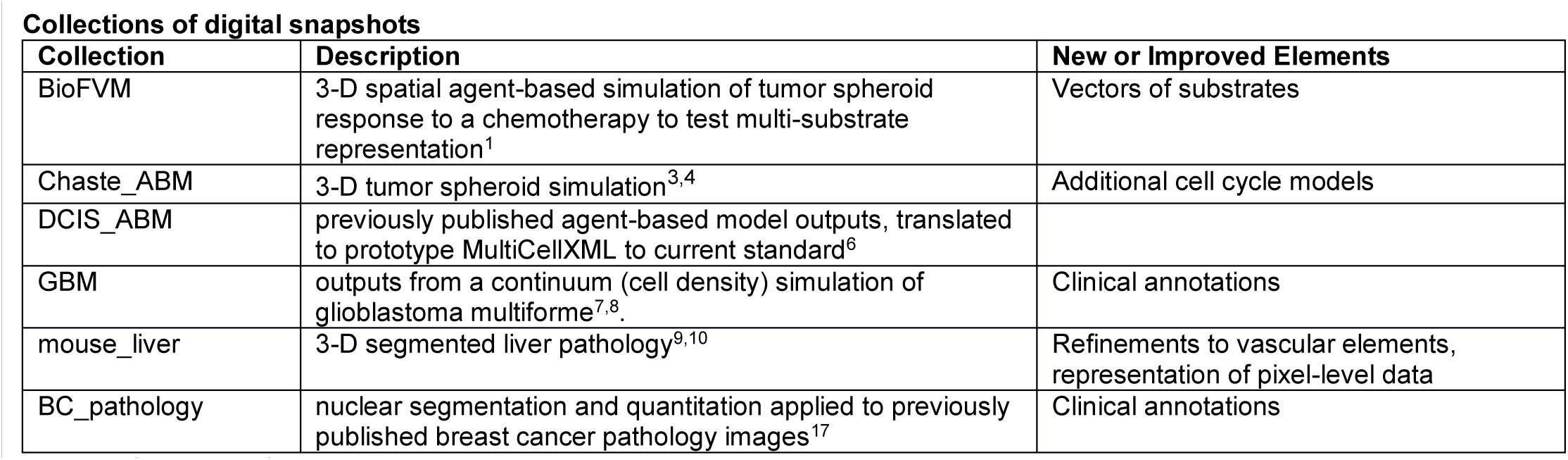
Summary of digital snapshots in the library.

Collections of digital snapshots
| Collection | Description | New or Improved Elements |
| --- | --- | --- |
| BioFVM | 3-D spatial agent-based simulation of tumor spheroid response to a chemotherapy to test multi-substrate representation<sup>1</sup> | Vectors of substrates |
| Chaste_ABM | 3-D tumor spheroid simulation<sup>3,4</sup> | Additional cell cycle models |
| DCIS_ABM | previously published agent-based model outputs, translated to prototype MultiCellXML to current standard<sup>6</sup> |  |
| GBM | outputs from a continuum (cell density) simulation of glioblastoma multiforme<sup>7,8</sup>. | Clinical annotations |
| mouse_liver | 3-D segmented liver pathology<sup>9,10</sup> | Refinements to vascular elements, representation of pixel-level data |
| BC_pathology | nuclear segmentation and quantitation applied to previously published breast cancer pathology images<sup>17</sup> | Clinical annotations |

While iterating on the digital snapshot standard, we recorded simulation data including: previously reported 2D agent-based ductal carcinoma in situ (DCIS) simulation data^6^ an agent-based simulation of a 3D tumour spheroid using Chaste^3,4^, 3D continuum simulations of vascularized brain cancer^7,8^ and lymphoma^28,29^, and a 2D simulation of vascularized tumour growth using BioFVM^1^. We also used digital snapshots to record nuclear morphometric properties on segmented H&E breast cancer pathology images^17^. Using the same data elements for experimental, clinical, and simulation data was a motivating factor for founding the MultiCellDS project, as it allows more direct comparison of simulation and clinical / experimental validation data, as well as more direct initialization of simulations and unified analysis platforms.

## Discussion

The MultiCellDS Project has grown from a single lab’s motivation to an international community, with the common goal of tackling the fundamental problem of recording and sharing multicellular data. By creating a standardized multicellular data representation with appropriate metadata, the research community can proceed to build big data repositories that are truly multiscale—including measurements from the molecular, cellular, multi-cellular, tissue, and whole-organism/patient scales. With a growing collection of standardized data, we can develop an ecosystem of stronger, mutually compatible software tools to quantitate and analyse data, devise new theories and predictions, and ultimately yield new discoveries that drive science and medicine.

Learning from notable prior efforts^70-72^ (see **Methods** as well), we focused on the core problem of representing and structuring data, rather than annotating mathematical models. To accelerate development, we adopted a “startup” development model with a small core team responsible for leading development, an invited multidisciplinary review panel to review, refine, and test the standard, and frequent engagement with the broader scientific community through public talks, conferences, and social media outreach. We employed a use case-based development cycle (choosing test problems and brainstorm solutions; discussing the prototype data elements; testing and refining until the tests succeed) in multiple rounds. In Round 1, we worked on describing microenvironment-dependent cell phenotype and cell line metadata. Round 2 focused on describing simulation and segmented pathology data, while Round 3 (ongoing) worked to describe clinical outcome metadata and collections of data. Where possible, we leveraged existing ontologies and standards where well-developed data elements existed; we developed new data elements or extended standards when necessary. See **Supplementary Material Timeline** for additional information.

MultiCellDS’ three main data types, digital cell lines (DCLs), digital snapshots, and collections, respectively allow a broad characterization of cell behaviour across many microenvironmental conditions, record spatial information (individual cells or cellular populations, distribution of microenvironmental substrates) at a single time, record key metadata, and organize the former two types into logical groupings (e.g., variants of MCF-7, time series data, data replicates). By utilizing the same data, we facilitate better exchange and comparison of data between quantitative modellers, data scientists, experimentalists, and clinicians. The use case-driven development created a library of over 200 DCLs, spanning prokaryotes and eukaryotes, epithelial and stromal cells, and patient-derived and “standard” cell lines. These are stored in a public repository, MultiCellDB, at http://MultiCellDS.org/MultiCellDB. The work also generated a collection of digitized, segmented, and spatially annotated breast pathology data (as digital snapshots) and simulation outputs (which can be used to benchmark future models). The hierarchical nature of MultiCellDS, while challenging to traditional database methods as the data are not flat (easily represented in a two-dimensional table), is critical to maintain its organization, searchability, and extensibility.

Having agreed on a data standard with an initial public data repository, we will now turn our focus to broader community involvement, to encourage data and software tool contributions as well as refinements to the standard. This is already partly accomplished through aggregated citations: contributors are credited for contributing new measurements or improved analyses, and original sources are always attributed. The momentum of the effort may also incentivize participation: as more data and more software tools become available using the standard, new developers will have incentive to also adopt the standard to make use of those data and tools. However, at this early stage, we envision that additional incentives may be needed, such as data hackathons with prizes (to encourage creation of tools), partnerships with granting agencies and journals (to encourage publication of supplementary data in standardized formats), and collaborations with instrument makers (to enable simple, direct writing of standards-compliant data). Moreover, the repositories and software need to be searchable, flexible, and easy to use, to prevent user frustration that can ultimately doom any emerging technology. The MultiCellDS community must involve user feedback and experts in human-computer interaction when developing and refining tools. The built-in support for custom data elements should ensure sufficient flexibility, and the biologic functional-driven hierarchy of data elements helps to ensure that the data are searchable.

As we conclude the technical challenges of writing and aggregating data into shared digital cell lines, we face new challenges in curation and quality control. How do we assess the quality of a new measurement^9^? When should a new measurement replace an existing measurement? Who is responsible for and granted authority to make such decisions? These open questions can be addressed by forming community standards for curation—a natural evolution of the current standards review panel. While prior efforts exist for other biological domains (e.g. the International Society for Biocuration), the multicellular community must create curatorial data requirements suited to its own domain. As this important work progresses, the community can collectively implement heuristics to help pre-screen new data, based on prior annotated knowledge. (e.g. “Cell radius for human breast cancer cells should be between 5-15 µm; raise a warning flag and any values outside that range.”) In the future, machine learning could not only flag bad measurements but identify typical causes (e.g., culturing without a key growth factor) and suggest solutions, thus improving experimental workflows.

In the future, we will continue developing the MultiCellDS data elements and metadata to sufficiently record data from emerging clinical trials and machine learning experiments in digital pathology and will extend the standard to HDF5^73^ (a widely-supported compressible hierarchical data format in the physical sciences). MultiCellDS can currently describe cell phenotype broadly across many microenvironmental conditions. Future work will add depth by integrating subcellular scales (e.g., metabolomics, receptor trafficking, and other “omics”) into the phenotype datasets, leveraging established standards such as SBML at those scales. Standardizations for the dynamics of pharmacodynamics and pharmacokinetics are needed to truly characterize high-content drug screening data and predict clinical response. We will work closely with the PharmML team to develop these data elements while harmonizing data elements across PharmML, SBML, and other standards. We will also work with leading open source computational modelling projects (particularly Chaste, CompuCell3D^55^, PhysiCell^74^, and TiSim^75^) to build software cross-compatibility by supporting the MultiCellDS data format; indeed, this work has already begun.

The new MultiCellDS standard and repository represent a start to sharing well-structured, machine-readable multicellular data. Now that we have tackled the issues of how to consistently record and structure data spanning many sources and types, it is time to work on building larger databases of multiscale, multicellular data, as has recently been highlighted by USA’s Vice President Biden’s “Cancer Moonshot” project^76^ and countless advocates for big data, open science, and reproducibility. With large repositories of consistently structured data, we can apply machine learning, mechanistic modelling, and other quantitative techniques to the big data to start answering big and significant questions: What is the state of our collective knowledge today? Where are the biggest gaps? Which cell and tissue parameters are most predictive to cell, tissue, and disease behaviour? And if those measurements are currently lacking, what technological advances are needed to obtain them? Through a truly systematic representation of biological knowledge, we can rationally prioritize research directions at national and global levels, and potentially get the best return for limited funding resources.

MultiCellDS gets us off the ground by providing a consistent means to record multicellular data. The repository helps to collect and share standardized data. The community now has the opportunity as well as the responsibility to make use of the data: to contribute more data; to mine the data for subtle patterns that drive new hypotheses; to test new hypotheses in computational, experimental, and clinical models; and to unlock new knowledge that drives scientific progress and yields new therapies and strategies to improve human health.

## Method

Originating as MultiCellXML (MultiCellular eXtensible Markup Language)^6^, MultiCellDS grew and was impacted by other standardization efforts. MultiCellDS focused on satisfying unmet needs in model-independent data descriptions—rather than model representation—to facilitate interchange of data between computational modellers, experimental biologists, and clinicians. We adopted a “startup” organizational structure to promote faster development of the data standard. We employed a use case-driven development process to iteratively assess and refine the standard while ensuring that it could succeed at its primary tasks of (1) representing and sharing microenvironment-dependent cell phenotype data while preserving provenance, and (2) sharing standardized, model-independent spatial multicellular state data from many types of simulations and segmented pathology images.

### Lessons from prior standardization efforts

To date, there have been successful instances of shared biological data repositories in molecular biology (e.g., TCGA^35^) and clinical medicine (e.g., some clinical trial databases^77^), where the data are relatively homogenous (many records with identical data elements) and *flat* (non-hierarchical). However, at the intermediate scales— cell phenotype and multicellular organization—efforts have been less successful. Encyclopaedias such as the Cancer Cell Line Encyclopedia (CCLE)^78^ have successfully collected useful molecular information, but have generally lacked phenotypic and multicellular data; the mapping from genotype to phenotype is not always straightforward, and demonstrating how cells interact with one another requires an understanding beyond single cells in isolation. Moreover, encyclopaedias have a tendency towards recording static data, discouraging active community contributions of improved measurements and analyses.

In a key advance towards community-driven databases, BioNumbers^7^ allows much more active user contributions and provides a generalized search engine. However, the data are not organized by cell type, there is no adaptable, hierarchical organization to help searches and comparisons by function, and there is no consistent reporting of microenvironmental metadata (e.g., oxygenation conditions, and growth medium used), even as these are known to impact cell phenotype^42,43,79^. Earlier data standards have similarly worked to describe cellular data (e.g., The Systems Biology Markup Language (SBML)^70,71^, CellML^72^), but they have largely focused on representing individual mathematical or computational models and their corresponding parameter values, rather than annotating model-independent cell phenotype. Moreover, these well-established standards have focused primarily on subcellular and cell-scale properties; only recently has SBML begun drafting specifications to represent multicellular models with its Dynamic and Spatial Processes packages, but the complexity of representing a wide variety of mathematical models has necessitated a long-term development process (e.g. SBML’s history^80^).

The development of ontologies (e.g. GO^62,63^) has helped improve recording results, but we focus on a key difference between an ontology and a data standard: ontologies are like dictionaries by defining terms, and data standards are like grammars that give a predictable structure to the terms. Both are necessary to communicate data fully and should drive the development of one another. The Cell Behavior Ontology (CBO)^64^ has helped steer the development of MultiCellDS, and we have submitted improvements to the CBO.

We chose the MultiCellDS organization model and development process based upon the lessons learned from these earlier projects. We focused on representing model-independent phenotype data to avoid the inherent complexity of representing all current and possible future mathematical models. This had the additional benefit of allowing the same data elements to represent experimental, clinical, or computational data. We iteratively tested and refined the emerging standard against a series of use cases stemming from simulation, experimental, and clinical science. This made development more concrete while ensuring that the final standard achieved its design goals. We adopted a “startup” organizational model, where a core group of “invested” participants (based at the University of Southern California) took responsibility for drafting and developing the standard. The core group recruited a multidisciplinary review panel of leading biologists, clinicians, modellers, and data scientists, each stakeholders in their own domain to refine and improve the draft standard while completing the use cases. Public presentations at scientific meetings and social media interactions were used to solicit and incorporate feedback from the broader scientific community.

### Creating the MultiCellDS standard

MultiCellDS originated as a way to save agent-based simulation data in a model of DCIS of the breast^6^. Keynote talks and discussions with modellers and experimentalists at the NCI-funded Physical Sciences in Oncology Network (PS-ON) 2013 annual meeting^81^ found the need to standardize data at the multicellular scale, for improved sharing of phenotype measurements, experimental data, and simulation outputs. To drive rapid progress towards a working standard, we used a two-stage strategy: in the first year of the project, we assembled a core team at USC to integrate early consensus opinions from the 2013 PS-ON annual meeting into the preceding MultiCellXML draft specification. By the end of this year, we had developed a working draft standard for DCLs and digital snapshots.

In Year 2, we expanded the development process to a select multidisciplinary panel of computational modellers, clinicians, data scientists, software developers, and biologists. Panel members were recruited from the PS-ON and the broader scientific and medical communities. Social media engagement and public presentations were used to recruit further review panellists. The full panel membership list is given in the **Supplementary Material**. The review was organized into three rounds: Round 1 (completed) focused on DCLs, and Round 2 (completed) focused on digital snapshots for simulation and experimental data. Round 3 (in progress) is focused on clinical, pathologic, and radiologic data elements. The core team and review panel determined that the data standard should be released in parts, rather than waiting for the completion of all three rounds. This was to give faster utility, help accelerate adoption, and ease the cost and effort of software implementation, by allowing parts of the standard to be implemented as they are completed.

Each review round consisted of three phases. The first phase (“brainstorming”) used one-on-one interactions with panel members and the broader public through video conferencing and seminar talks to explore and propose data elements and their hierarchical relationships. XML schema files were maintained throughout this process to fully capture the definitions and allowed relationships in MultiCellDS XML files. The second phase (“formal review”) conducted a series of “town hall” style videoconferences with the review panel, to allow frank discussion of the elements and solicit feedback. The core group introduced refinements to the XML schema files between each town hall meeting. The third phase (“test-based refinements”) tested the new data elements against specific task(s) by the review panel members, to expose bugs and omissions in the standard. Refinements were made to the schemas until the tests could be successfully completed. See the timeline in the Supplementary Materials.

The Round 1 tests used clinical, experimental, and literature data to create DCLs for mammalian cells (cancer, stromal, and other types), non-mammalian cells (yeast), and prokaryotic cells (*S. aureus* and *E. coli*), with varying levels of detail (see **Results** and **Supplementary Material**.) Round 2 tests focused on representing simulation outputs for many discrete^3,4,6,53-56,82-84^ and continuum^28,51,85-87^ computational models and segmented nuclei in clinical pathology data. Round 3 tests will represent more general clinical pathology data.

### Use case: Building a library of digital cell lines

We have created DCLs for a wide variety of cell types, with the largest number of DCLs coming from (1) individual patients in cancer studies and (2) many cancer types from data provided by ATCC to the PS-ON^5^. We also created DCLs for Human Umbilical Vein Endothelial Cells (HUVECs), lymphoma, yeast and bacteria. We put these DCLs into an online searchable repository (MultiCellDB). We have found that the hierarchical nature of the DCLs was challenging to traditional database methods as DCL data are not flat (easily representable in a two dimensional table); this hierarchical nature is critical to maintain the organization, searchability, and extensibility of MultiCellDS. Each collection of DCLs presented unique annotation challenges, thus driving refinements in the data standard.

#### Systemizing data for previous PS-ON investigation

Earlier work^2^ measured a wide variety of cellular properties of MCF-10A and MDA-MB-231 cells. However, obtaining numerical information from this paper presented a set of challenges, since the data are not easily accessible. Data values are presented in plots, sometimes in the text of the paper or supplementary material, sometimes in a Word file contained in a compressed file stored on an unreferenced Wiki, and sometimes in Word documents that are not available online in any form at all (except by request from individual co-authors). We also encountered inconsistent formats and missing growth parameters. While presenting data in these various formats can speed up initial data exchange between collaborators (at the level of files, tables, and plots), it does not contribute to detailed information exchange (e.g., direct comparison and combination of measurements between groups), nor facilitate quantitative communication between computational models, other software packages, and databases. When creating the MCF-10A and MDA-MB-231 DCLs, we systemized the reported data and collected missing primary information from the authors to complete the DCLs. This systematization of the data thus continues the impetus for the original study. To successfully complete this set of DCLs, we created “dimensions” of data to record additional results. Often, a parameter is represented by a single value, but will be representative of a larger collection of non-temporal data points; for example, Young’s modulus measurements as the slopes of linear fits to multiple indentation measurements. We record this larger collection to retain the original knowledge and provide better confidence in future results; see Anscombe’s Quartet^88^.

#### Creating digital cell lines from experimental protocol information

The Physical Sciences in Oncology Bioresource Core Facility offers cell line information and standard operating procedures (SOPs) for 38 human cell lines derived from 7 organs, as freely-available PDF files^5^. We extracted a growth curve (as an image) from the PDF documentation for each cell line, processed the image in MATLAB to extract the cell counts and standard deviations at several time points, and performed a nonlinear least squares fit of the extracted data to exponential and logistic growth models (see the Supplementary Materials) to extract cell birth rate parameters. (The software to reproduce this work can be found at https://source-forge.net/projects/multicellds.) We also extracted metadata from the SOPs, and combined these with the cell birth rate parameters to create 38 minimal DCLs which can later be extended by the broader community, analogously to Wikipedia “stubs”^89^. Work on this early collection of DCLs (with its variety of cancer types and cellular origins) exposed the need for detailed biological cell line metadata elements. We introduced metadata related to a cell’s origin, so that we could distinguish between the different types of cancers (e.g. breast cancer vs. colon cancer). This logically extended to have other subelements, e.g. disease or species/organisms.

#### Two collections of patient-derived digital cancer cell lines

Patient cohorts permit studying the variability of cell phenotype, albeit over a larger range of genetic variability. The addition of clinical data allows a systematic way of relating various phenotype measurements to clinical outcomes. We refined the MultiCellDS draft standard to record phenotype measurements for two collections of patient cell lines: glioblastoma multiforme (GBM), based upon radiology, clinical outcome, and limited pathology data^10^, and DCIS of the breast, based upon morphometric and other pathology-based measurements^18^. The GBM collection includes 39 patients with motility data and an additional 133 patients with partial data, but not as complete as the other patients^10^. To successfully create the GBM DCLs, we introduced (de-identified) patient- and disease-focused metadata to the standard. For the DCIS collection, we created 17 patient-derived DCLs, using an updated calibration protocol which improved upon previous methods^6^; see the Supplementary Materials. To create these DCLs, we required new metadata elements to describe the patient clinical staging information, as well as organism, organ, and tissue identifying elements.

#### Other digital cell lines

The lymphoma DCLs stem from two papers^28,29^, and include geometrical properties, PKPD responses, and cell cycle data. To ensure that both eukaryotic and prokaryotic cells can be successfully represented, we created DCLs for *Staphylococcus aureus (S. aureus)* and *Escherichia coli* (*E. coli*) bacteria to record proliferation, size, and mechanics data^26,27^. Future MultiCellDS releases will include new data elements for bacterial biofilms, as these properties are known to impact cell proliferation, death, and response to antibiotics and other environmental stressors^90,91^. Human Umbilical Vein Endothelial Cells (HUVECs) have several different behavioural “states”: as tip, stalk, and phalanx cells^22^, and we denote the different phenotype datasets using keywords. The HUVEC DCLs recorded these proliferation parameters, transport processes, motility, size and motility data elements based previous work^16,21,92,93^. Data reported in nine different papers had parameters that could have potentially been used for DCLs^21,92,94-107^; however, these papers reported qualitative data (e.g., an environmental factor increases motility, but with unspecified magnitude) or relative quantitative data (e.g., a proliferation rate may double relative to control, but with no scale reported). This makes it difficult to combine and compare data from multiple sources.

The yeast *Saccharomyces cerevisiae* is a model organism for eukaryotic cells and cell cycling. Thus yeast are an important use case for DCLs, for building predictive mathematical models as well as translating observed microscopic behaviour to macroscopic population level behaviour. Because yeast can be grown in liquid or solid media, this DCL details the solid media preparation^23^. The replicative behaviour of yeast has been quantified in many studies^24^. We note that the cell cycle of yeast is different for cells in their first division, which have an extended G_1_ phase. The replicative life span of yeast is thought to depend on the number of cell divisions rather than a specific time span; the average life span is around 25 divisions^25^. Yeast have cell walls rather than cell membranes – as in human cells – and the mechanical properties of these walls have been documented^108^. Finally, yeast cells reproduce through a budding mechanism where newly born daughter cells do not inherit the replicative age of the mother. Thus, the yeast use case required us to introduce new data elements to record a generation count that increases with each division.

## Supplementary material

The MultiCellDS project website is hosted at http://MultiCellDS.org. A list of MultiCellDB portals, including the MultiCellDB reference repository, can be found at http://portals.MultiCellDS.org.

1. User-focused overview of the data standard. [https://dx.doi.org/10.6084/m9.figshare.4269254.v1].
2. List of supported cell cycle representations. [https://dx.doi.org/10.6084/m9.figshare.4269263.v1]
3. Computer-generated documentation on the full standard, based upon the XML schema. [https://dx.doi.org/10.6084/m9.figshare.4269269].
4. The XML schema that official encodes the data standard. [https://dx.doi.org/10.6084/m9.figshare.4269272.v1 and https://dx.doi.org/10.6084/m9.figshare.4269275].
5. OWL ontology. [http://MultiCellDS.org/ont/multicellds.owl].
6. A protocol to transform DCIS pathology data into patient-derived digital cell lines. [https://dx.doi.org/10.6084/m9.figshare.4269248.v1].
7. Mathematical models used in the Chaste demonstration of MultiCellDS digital snapshots [https://dx.doi.org/10.6084/m9.figshare.4272242].
8. Matlab script used to help create ATCC-based digital cell lines [https://sourceforge.net/projects/multicellds/files/Tools/ATCC_to_digital_cell_lines/]
9. Community norms for curation, versioning, and new data elements [https://dx.doi.org/10.6084/m9.figshare.4272374.v1].
10. Current MultiCellDS invited reviewers. [http://MultiCellDS.org/Team.php#review_panel]
11. MultiCellDS invited reviewer (rounds 1-3, through Nov. 2016). [https://dx.doi.org/10.6084/m9.figshare.4272197]

All MultiCellDS documentation can be found at http://MultiCellDS.org/Documentation.php.

A list of MultiCellDS-compatible software is maintained at http://software.MultiCellDS.org.

